# A simplified two-plasmid system for orthogonal control of mammalian gene expression using light-activated CRISPR effector

**DOI:** 10.1101/2024.12.13.628044

**Authors:** Shruthi S. Garimella, Shiaki A. Minami, Anusha N. Khanchandani, Susannah R. Schaffer, Priya S. Shah

## Abstract

**Background:** Optogenetic systems use light-responsive proteins to control gene expression with the “flip of a switch”. One such tool is the <u>l</u>ight <u>a</u>ctivated <u>C</u>RISPR <u>e</u>ffector (LACE) system. Its ability to regulate gene expression in a tunable, reversible, and spatially resolved manner makes it attractive for many applications. However, LACE relies on delivery of four separate components on individual plasmids, which can limit its use. Here, we optimize LACE to reduce the number of plasmids needed to deliver all four components.

**Results:** The two-plasmid LACE (2pLACE) system combines the four components of the original LACE system into two plasmids. Following construction, the behavior of 2pLACE was rigorously tested using optogenetic control of enhanced green fluorescent protein (eGFP) expression as a reporter. We optimized the ratio of the two plasmids, measured activation as a function of light intensity, and determined the frequency of the light to activate the maximum fluorescence. Overall, the 2pLACE system showed a similar dynamic range, tunability, and activation kinetics as the original four plasmid LACE (4pLACE) system. Interestingly, 2pLACE also had less variability in activation signal compared to 4pLACE.

**Conclusions:** This simplified system for optogenetics will be more amenable to biotechnology applications where variability needs to be minimized. By optimizing the LACE system to use fewer plasmids, 2pLACE becomes a flexible tool in multiple research applications.

## Background

Inducible mammalian gene expression systems are valuable for research, biomedical, and biotechnology applications. In research applications, inducible gene expression can uncover the impacts of a specific gene with respect to their biological system or be used for biosensing [1]. For biomedical applications, inducible gene expression can be used to improve tissue engineering efficiencies by controlling expression of stem cell multipotency, proliferation, or differentiation factors [2]. Inducible gene expression can also be used to improve biosynthetic pathway activity [3, 4] or control protein expression when cell density is optimal to maximize productivity [5, 6].

Orthogonal modes to control gene expression can be especially useful depending on the application and can even be multiplexed. However, each system is well suited for different applications. Chemically induced systems (e.g. doxycycline, cumate, rapamycin) are by far the most prevalent and easy to implement. However, the chemical inducers can have off-target effects, require dilution or media replacement for reversal, and lack spatial control [1, 7]. Temperature-based systems use heat or cold shock to stimulate the system [8]. This is simple to use and reverse, but still lacks spatial control. Light-inducible or optogenetic systems function by using a specific wavelength of light to excite a light-activated protein engineered to control biological process [9–12]. Optogenetic systems are uniquely suited for reversible and precise spatial control of gene expression because the light can easily be switched off and targeted to specific location via optical setups [13–16].

Several optogenetic systems have been developed recently using actuation by different wavelengths, including UV, blue, green, and red spectra [15, 17–21]. Red light systems may be of interest from an optical standpoint, since the lower energy light is less cytotoxic. In cells, red light also has lower absorption and scattering coefficients compared to blue light. Consequently, red light provides higher spatial control and deeper penetration into thicker samples [21, 22]. The iLight and iLight2 systems take advantage of a conformational change in a red light-responsive protein to expose a DNA binding domain and drive gene expression [23–25]. However, the system relies on an engineered DNA binding domain that can only bind to a synthetic promoter. Thus, iLight cannot be used to control endogenous genes without re-engineering the DNA binding domain or through genetic modifications to the gene expression regulatory circuit of the target gene.

Blue-light systems are the most widely used optogenetic systems because of their higher chromophore efficiencies and the development of several light-responsive systems engineered to have improved performance [26, 27]. Light-Activated <u>C</u>RISPR <u>E</u>ffector (LACE) is one such system that takes advantage of protein dimerization following excitation by blue light. The light-responsive protein CRY2 is fused to viral protein 64 (VP64), a transcriptional activation domain derived from herpesvirus. CIBN is fused to deactivated Cas9 (dCas9), which is targeted to a minimal CMV promoter or an endogenous promoter using an appropriate guide RNA (gRNA). CRY2 undergoes a conformational change when stimulated by blue light and dimerizes with CIBN. CRY2-CIBN dimerization brings VP64 to the promoter to activate gene expression. As a reversible system, gene expression turns off when light is removed, when CRY2 dissociates from CIBN [18, 28].

LACE is a powerful tool that enables modular optogenetics with flexibility to target exogenous and endogenous genes. Endogenous gene expression can be controlled if a suitable gRNA can be designed [18]. However, the LACE system requires delivery of four plasmids, which may limit how many cells receive all four components, especially in hard-to-transduce cell types. This limitation can increase variability in the cellular response, and increase the complexity of applying LACE to cell types relevant for biomedical and biotechnology applications [28]. Here, we improve upon the LACE system by decreasing the number of plasmids required and re-optimizing the ratio of the two combined plasmids. We compare two-plasmid LACE (2pLACE) to four-plasmid LACE (4pLACE) and find that 2pLACE has similar dynamic range, tunability, and kinetics to 4pLACE with the added advantage of being more consistent. Thus, 2pLACE provides improved outcomes for optogenetic applications.

## Results

### Design and plasmid ratio optimization of the 2pLACE system

We first set out to design a LACE system that would be easier to use and more efficient for various applications. The LACE system requires four different components to be delivered to cells. Originally, these four components were delivered on four separate plasmids by transient transfection [18, 28]. Consequently, the upper limit on LACE activation is dictated by the cells that receive sufficient amounts of all four plasmids. We therefore hypothesized that reducing the number of plasmids used to deliver LACE components would improve LACE behavior. We therefore designed a system to reduce the number of plasmids from four to two by combining the CRY2-VP64 and minCMV-eGFP components and combining the CIBN-dCas9 and gRNA components (Figure 1 and S1). These plasmids were synthesized commercially based on our design using a pCDNA3.1 backbone.

**Figure 1.**
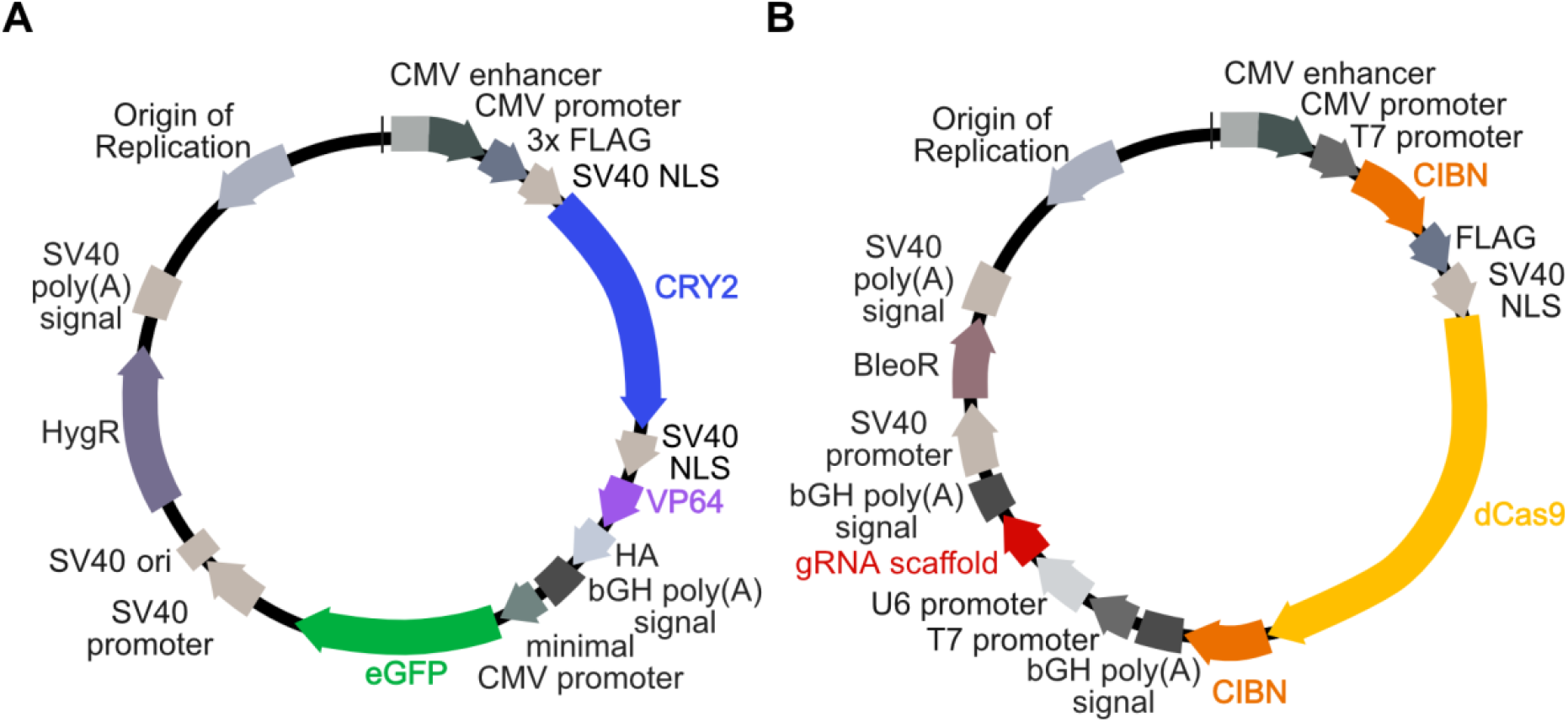
Schematic of 2pLACE design. (A) Expression cassettes for CRY2-VP64 and target gene (eGFP) with minimal CMV promoter and gRNA binding sites were combined into a single plasmid (CRY2-eGFP). (B) CIBN-dCas9-CIBN and gRNA expression cassettes were combined into a second plasmid (CIBN-gRNA). Each plasmid was designed to include an SV40 origin of replication (ori) for transient plasmid maintenance in cells expressing Large T-antigen. A distinct selection marker in each plasmid (Hygromycin and Bleocin) was included to enable stable cell line generation.

As the ratios of the LACE components may influence leaky gene expression, maximal activation, and the dynamic range, we optimized the transfection conditions by varying the mass ratios of the two plasmids. Cells were kept in the dark or activated with pulsing blue light for 24 hours, and eGFP reporter fluorescence was analyzed through flow cytometry. As the ratio of CRY2-eGFP to CIBN-gRNA plasmid was increased, the background expression of eGFP in the dark consistently increased. Expression of eGFP following activation also increased as a function of plasmid ratio, but peaks at a ratio of 3:7 and slowly decreased beyond that (Figure 2A and S2). Dynamic range (ratio of light:dark signal) started high and consistently decreased as the CRY2-eGFP to CIBN-gRNA ratio increases (Figure 2B). To balance high eGFP expression following light activation and high dynamic range, the optimal ratio of 3:7 was used for all subsequent experiments.

**Figure 2.**
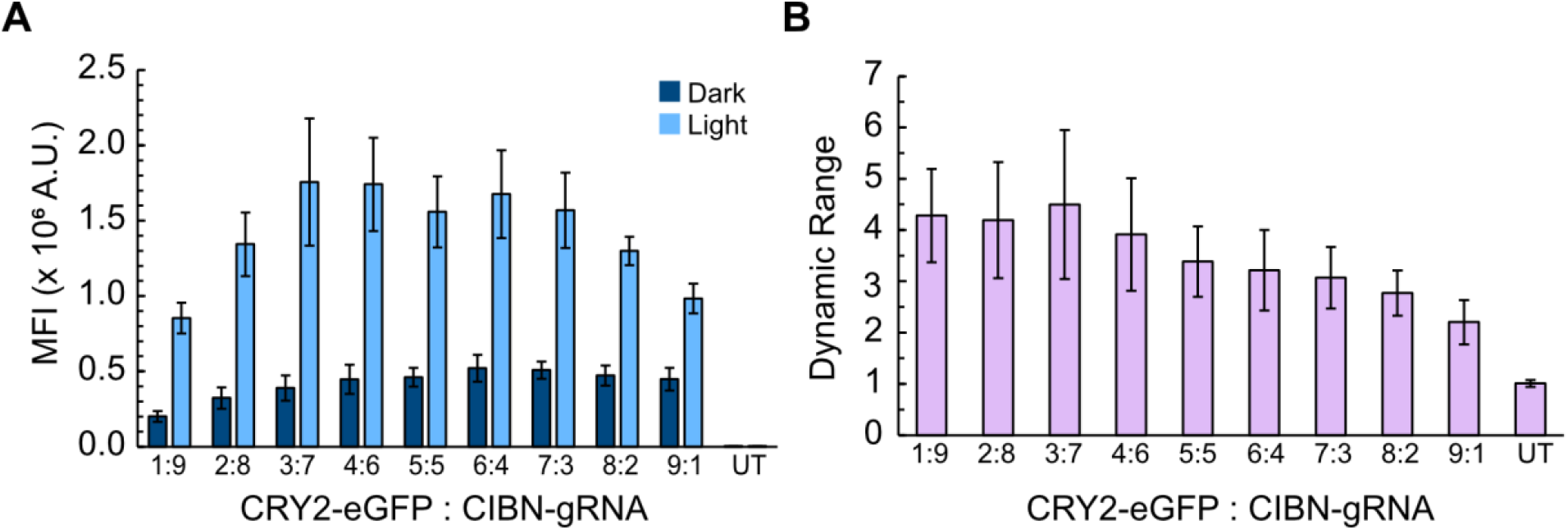
2pLACE plasmid ratio optimization for high absolute activation and dynamic range. (A) Mean fluorescence intensity (MFI) was measured for cells transfected with an increasing ratio of CRY2-eGFP:CIBN-gRNA plasmids followed by 24 hours of incubation in light or dark. (B) Dynamic range for different plasmid ratios from (A) was calculated as MFI in light vs MFI in dark. Plots represent the mean of three independent biological replicates and error bars represent standard deviation. UT (untransfected) is the negative control.

### Characterization of transient 2pLACE activation dynamics

Next, we measured the responses of the system to lighting conditions to assess tunability. We first tested the sensitivity of the system to LED intensity, where intensity was varied and eGFP fluorescence was measured 24 hours post-activation. Experiments were performed using the optoPlate to enable high-throughput measurements in a 96-well format [29]. We found that the tunable range of eGFP expression occurred at intensities between 0-2 mW/cm^2^, with saturation behavior beyond those intensities (Figure 3A). Significant activation occurred with light intensity as low as 0.12 mW/cm^2^, which was the lowest intensity tested (Figure S3). However, the activation was still quite variable for low intensities of light. For subsequent experiments, we used an intensity of 9.23 mW/cm^2^. This corresponds to the intensity used to test the optimal mass ratio of 2pLACE (Figure 2) as well as the intensity used in previous work [28]. Maintaining a similar intensity as the 4pLACE experiments also allows for direct comparison of system response.

**Figure 3.**
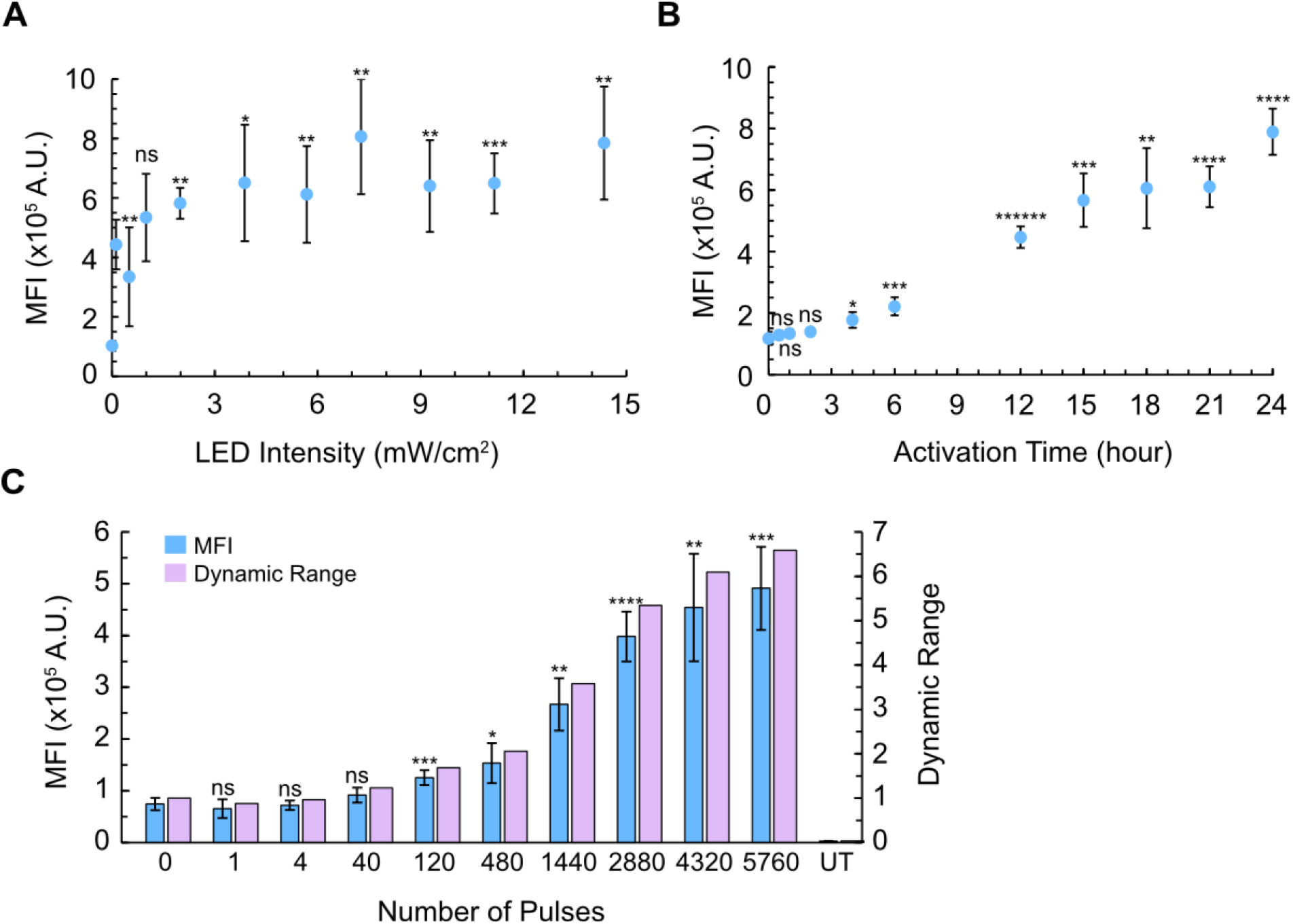
Characterization of 2pLACE activation dynamics. (A) eGFP MFI as a function of LED intensity. (B) eGFP MFI as a function of activation time. (C) eGFP MFI and dynamic range as a function of the number of LED pulses applied. UT (untransfected) is the negative control. Plots represent the mean of six technical replicates and error bars represent standard deviation. MFI significance was calculated with a one-way ANOVA and Tamhane’s T2 post-hoc test to the dark state condition with 0 hours of light activation or 0 pulses. * p < 0.05, ** p < 0.01, *** p < 0.001, **** p < 0.0001, ***** p < 0.00001 ****** p < 0.000001, and ns is not significant.

We next measured activation kinetics. We activated cells transfected with 2pLACE for increasing amounts of time using the same pulse frequency. eGFP fluorescence was measured by flow cytometry. This timecourse experiment showed that the system had an initial lag in expression before reaching significant but minimal expression as early as 4 hours. Activation continued to increase with prolonged activation time (Figure 3B). The activation kinetics of 2pLACE confirmed that short exposure or activation times turns the system on, but the longer the system is exposed to light, the greater the protein can be expressed.

Our kinetics experiments suggest that prolonged exposure to light maximizes gene expression. However, blue light can be cytotoxic [30]. Moreover, some applications may benefit if the system can be activated with a short exposure to light [30–32]. We therefore evaluated the sensitivity of the system to the total number of pulses to determine if there was a critical threshold of pulses for efficient activation while still allowing time after activation for the protein to be expressed. Cells were transfected with 2pLACE and activated with blue light for the specified number of pulses and the same pulse frequency. eGFP fluorescence was measured at 24 hours after pulsing was initiated (Figure 3C). Significant activation occurred at 120 pulses, which corresponds to 0.5 hours of activation time. Interestingly, this is lower than the activation time required for significant activation in our kinetics experiment (Figure 3B) and suggests that there is a lag between CRY2 association with the promoter and the accumulation of a detectable amount of protein in this system. However, even though significant activation was observed for 0.5 hours of pulsing, the reporter signal only reached substantial values starting at 1440 pulses, corresponding to 6 hours of activation time. Absolute activation and dynamic range continued to increase to 2880 pulses, which corresponds to 12 hours of activation time. Thus, 2pLACE can activate gene expression with as little as 0.5 hours of activation; however there are benefits to prolonged activation.

### Comparison of 2pLACE activation to 4pLACE

By reducing the number of plasmids needed, the 2pLACE system offers more ease of use compared to 4pLACE. To this point, we directly compared the light activation characteristics of both systems. We measured eGFP fluorescence by flow cytometry with and without 24 hours of light activation to understand the maximum expression of both systems (Figure 4). 2pLACE achieves a similar magnitude of expression as 4pLACE, and notably with less variability (Figure 4A). While there is no significant difference between the on states of the two systems, the consistency of 2pLACE is an improvement if the system is to be applied. We also compared activation kinetics of 2pLACE and 4pLACE over a 24 hour period. Both systems activated with similar kinetic behavior (Figure 4B). The matching activation profiles indicate the minimal system does not alter the dynamics of the LACE components. Thus, 2pLACE offers a similar activation profile to 4pLACE with improved consistency in activation.

**Figure 4.**
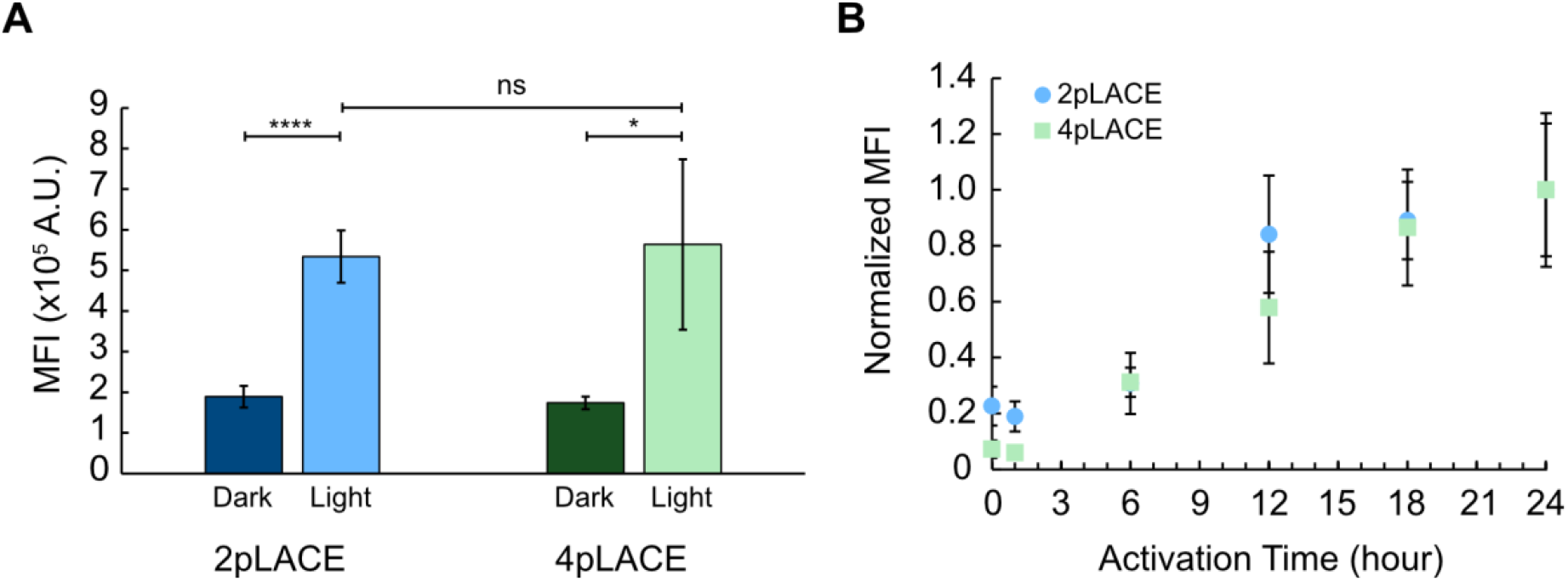
Comparison of 2pLACE and 4pLACE activation kinetics. (A) eGFP MFI was measured for cells transfected with 2pLACE and 4pLACE 24 hours post activation with and without blue light. (B) Activation behavior of 2pLACE and 4pLACE over 24 hours of activation. Data is normalized to the maximum activation of cells. Plots represent the mean of six technical replicates and error bars represent standard deviation. Significance was calculated with a one-way ANOVA and Tamhane’s T2 post-hoc test. * p < 0.05, **** p < 0.0001, and ns is not significant.

## Discussion

Optogenetic systems such as LACE are novel systems for inducible gene expression and have the potential to overcome limitations such as off-target effects, reversibility, cost, and spatial control associated with other inducible systems. These advantages make LACE a powerful and flexible tool for a variety of applications, provided all four LACE components can be introduced to target cells. However, the original LACE system was encoded on four different plasmids, limiting its potential use cases.

Here, we optimized the LACE system by reducing the number of plasmids needed to deliver the key components from four to two plasmids (Figure 1). We varied plasmid stoichiometry and found that a 3:7 plasmid ratio optimizes for absolute activation and dynamic range (Figure 2). The system can also be tuned using different blue light intensities, where significant activation occurs at 0.12 mW/cm^2^ and saturates after 2 mW/cm^2^ (Figure 3A). A kinetics time-course shows that the system is activated by 4 hours but improves with increasing activation time (Figure 3B). Interestingly, when a short period of activation is followed by a prolonged period for gene expression products to accumulate, significant expression is achieved after only 0.5 hours of light activation (Figure 3C). This suggests there is a delay between complex formation at the promoter and translation. These results also reiterate that activation occurs on short timescales, but prolonged activation results in higher gene expression.

Comparing 2pLACE and 4pLACE directly showed that the systems have similar absolute activation and dynamic range after a maximum 24 hours of activation; however the 2pLACE system provided a more consistent signal (Figure 4A). We speculate that the improved consistency of 2pLACE comes from reduced variability in cells receiving sufficient amounts of all four LACE components. Reducing the number of plasmids from four to two also does not alter the activation kinetics (Figure 4B). We hypothesize that many of the metrics between 2pLACE and 4pLACE are similar because the components have not changed. However, the reduced variability and the ease of use give 2pLACE improved potential for various synthetic biology applications.

Optogenetic systems like 2pLACE are low cost inducible systems ideal for use in low resource environments. The implementation of optogenetics in industrial applications can ease the high costs of serums and antibiotics that are common to improve yield in bioreactors and activate chemically induced systems [32]. For processes that require temporal control of gene expression, optogenetics also avoid costly media changes. 2pLACE’s ease-of-use means it can be applied to hard-to-transfect and more biologically relevant cells lines, such as primary cells used in tissue engineering for regenerative medicine and cultivated meat [33]. Cultivated meat is a growing area of interest as a sustainable solution for improving food accessibility [34, 35]. However, cost is critical to the industry’s viability. Easier-to-use optogenetic tools like 2pLACE could help bend the cost curve for the cultivated meat industry. For example, muscle stem cells benefit from expression of proliferative proteins for scale up, but these same proteins prevent differentiation into muscle cells [36]. Temporal control of proliferative gene expression by 2pLACE could be used to strategically promote proliferation during scale up. Differentiation could proceed unimpeded with a flip of the switch to turn off blue light activation of proliferative gene expression.

By optimizing the number of plasmids needed, 2pLACE shows more consistent behavior that improves its applicability as a tool. Yet, to be broadly applicable, additional improvements are still needed. Applying 2pLACE in industrial biotechnology settings will require stable expression of components to ensure activation over longer timescales. The work presented here characterizes 2pLACE under transient transfection conditions and are not necessarily compatible with long timescale processes. Our previous efforts to create a stable expression system for LACE using lentiviral vectors were hampered due to leaky expression in the dark state, and an inability to efficiently select cells that received all four components [28]. To this effect, we designed the 2pLACE system to have antibiotic resistance genes (Hygromycin and Bleocin, Figure 1) as distinct selection markers for each plasmid. Again, having fewer plasmids to encode all four components has advantages since fewer selection markers are required for stable cell line engineering. Stable expression of 2pLACE with prolonged activation by blue light could also lead to phototoxicity to the cells over long timescale scenarios [30]. Using a lower intensity or pulse frequency of blue light could overcome this toxicity.

The simplicity and versatility of 2pLACE makes it attractive for many applications, but it will still require specialized setups. For large-scale industrial applications, specific photobioreactors would need to be designed or current bioreactors would need to be outfitted with LEDs. We previously showed that blue light can propagate through a bioreactor to activate cells, but scattering is a major limitation that can dictate the upper limit on reactor size [32]. Specialized reactor designs with LEDs embedded throughout the reactor could also overcome the limitations associated with light propagation, though that may be costly. Ultimately, specialized smaller-scale applications like just-in-time or personalized medicine biomanufacturing may be more fruitful applications of 2pLACE.

## Conclusions

In conclusion, LACE is a reversible and tunable optogenetic tool that can be designed to target a specific gene of interest given an application. By reducing the number of plasmids needed to deliver all necessary components, 2pLACE becomes a flexible tool for synthetic biology and bioproduction applications. The use of two plasmids ensures consistent signal while maintaining reversibility and tunability of the LACE system. While the requirement of blue light can limit its applicability in certain applications, 2pLACE broadens the accessibility of inducible gene expression tools for low-resource environments and biologically relevant models.

## Methods

### Plasmids

The 2pLACE plasmids, pcDNA3.1-Cry-GFP-Hygro and pcDNA3.1-CibN-gRNA-Zeo, were commercially synthesized (Genscript). Full plasmid maps are included as supplementary files. 4pLACE plasmids, pcDNA3.1-CRY2FL-VP64, pcDNA3.1-CibN-dCas9-CibN, pGL3-Basic-8x-gRNA-eGFP, and gRNA-eGFP-Reporter, were gifted from Charles Gersbach (Addgene plasmid #60554, 60553, 60718, and 60719, respectively).

### Cell Culture

HEK293T cells (ATCC-No. CRL-11268) were cultured in DMEM (Gibco) supplemented with 10% FBS (Gibco). Cells were cultured in a humidified incubator at 37°C and 5% CO_2_. Cell density and viability were measured prior to cell plating using trypan blue and an automated cell counter (TC20, Bio-Rad). Cells were tested monthly for mycoplasma by MycoStrip (Invivogen) or PCR with the primers targeting 16S rRNA (Table 1).

**Table 1.**
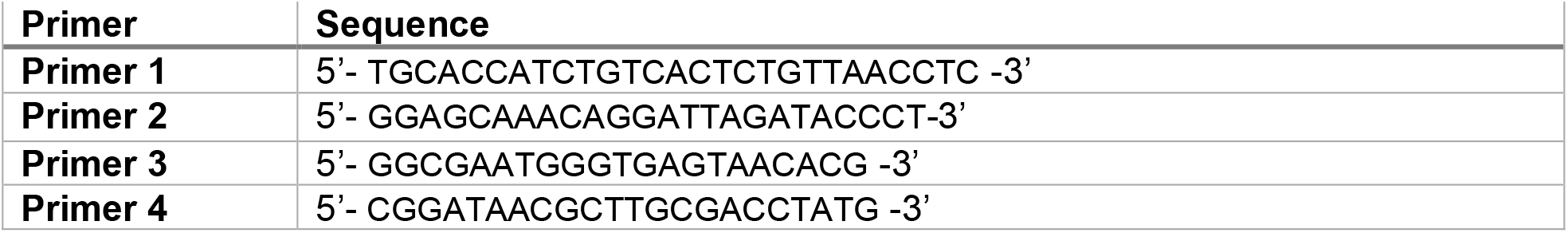
Primer sequences for mycoplasma pCR

### Transient transfection

Twenty-four hours before transfection, cells were seeded in a 96 well #1.5H glass bottom plate (Cellvis) at a density of 0.035 x 10^6^ cells per well. Each well was transfected with a total of 100 ng of DNA. The plasmid ratios for 2pLACE were relative proportions. Plasmid ratios for 4pLACE were calculated based on mass and molecular weight of plasmids relative to gRNA-eGFP-Reporter (Table 2).

**Table 2.** Relative mass ratios of plasmids transiently transfected by LACE system

| Plasmid Name | Relative Mass Proportions | Mass per well (ng) | System |
| --- | --- | --- | --- |
| **pcDNA3.1-Cry-GFP-Hygro** | 3 | 30 | 2pLACE |
| **pcDNA3.1-CibN-gRNA-Zeo** | 7 | 70 | 2pLACE |
| **pcDNA3.1-CRY2FL-VP64** | 2.06 | 20.6 | 4pLACE |
| **pcDNA3.1-CibN-dCas9-CibN** | 2.94 | 29.4 | 4pLACE |
| **pGL3-Basic-8x-gRNA-eGFP** | 4 | 40 | 4pLACE |
| **gRNA-eGFP-Reporter** | 1 | 10 | 4pLACE |

Transfections used PolyJet DNA In Vitro Transfection Reagent (SignaGen) with a PolyJet:DNA volume to mass ratio of 3:1 according to manufacturer instructions.

### Light activation

Cells were activated with LEDs at 465 nm twenty-four hours post transfection. For the optimization of plasmid ratios, a breadboard with 8 mm LEDs at a pulse frequency of 0.067 Hz. For all other experiments, light activation was performed using an optoPlate-96 [29]. LED intensities and pulse frequencies were regulated with an Arduino Micro microcontroller. For experiments performed in 6-well plates, samples in the dark were wrapped in aluminum foil to ensure isolation from light. LED intensities were measured using the 1931-C Newport optical power meter.

### Flow cytometry

Under red light, cells were trypsinized (0.05% Trypsin-EDTA, Gibco) and resuspended in cold DPBS (Gibco) with 1% FBS (FACS buffer). After centrifugation at 4°C for 10 minutes at 1000 RPM, 80% of the supernatant volume was removed and pellets were resuspended in 250 µL cold FACS buffer. Samples were wrapped in aluminum foil to ensure isolation from light following preparation for flow cytometry. A 488 nm laser in a CytoFLEX S Flow Cytometer was used for all analytical flow cytometry experiments. A target value of 10,000 total events was reached for every sample before gating for live, single, fluorescing cells. Cells were gated at an intensity of 5×10^3^ on the FITC-GFP channel. This corresponded to ∼0.5% of cells falling into this gate for an untransfected negative control.

### Statistical Analysis

Statistical analysis was done using a one-way ANOVA test with Tamhane’s T2 post-hoc test for its conservative analysis because it assumes unequal variances. Python packages scipy.stats f_oneway and scikit_posthocs were used for the ANOVA and post-hoc test respectively.

For the biological replicates, standard deviation was calculated with error propagated:

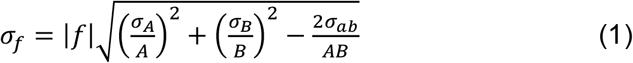

*σ*_*f*_ is the standard deviation of the dynamic range, *f* is the dynamic range value, *A* is the MFI value for a light activated sample, *B* is the MFI value for the dark state sample, *σ*_*A*_ and *σ*_*B*_ are their standard deviations, respectively. *σ*_*AB*_ is the covariance of the light and dark conditions. We assumed that the two conditions are not correlated, therefore the covariance is 0.

## List of Abbreviations

LACE: Light activated CRISPR effector
CRISPR: Clustered Regularly Interspaced Short Palindromic Repeats
dCas9: dead Cas9
gRNA: Guide RNA
eGFP: Enhanced green fluorescent protein
MFI: Mean fluorescence intensity

## Declarations

### Ethics approval and consent to participate

No ethics approval or consent was required for this research.

### Consent for publication

No consent for publication was required for this research.

### Availability of data and materials

The processed datasets supporting the conclusions in this article are included within the article. All raw data used to generate plots and materials will be made available upon request.

## Authors’ contributions

SSG, SAM, and PSS conceived of the project. All authors designed experiments. SSG, SAM, ANK and SRS performed experiments and analyzed data. SSG, SAM, ANK, and PSS wrote and edited the manuscript. All authors read and approved the final manuscript.

## Acknowledgements

We thank Bridget McLaughlin, Jonathan Van Dyke, and Ashley Karajeh for their technical assistance on flow cytometry experiments. We thank the Moule Lab at UC Davis for the use of the 1931-C Newport optical power meter.

## Supporting Information

See attached document.

