## Supplementary Material for "A simplified two-plasmid system for orthogonal control of mammalian gene expression using light-activated CRISPR effector"

### Supporting Information

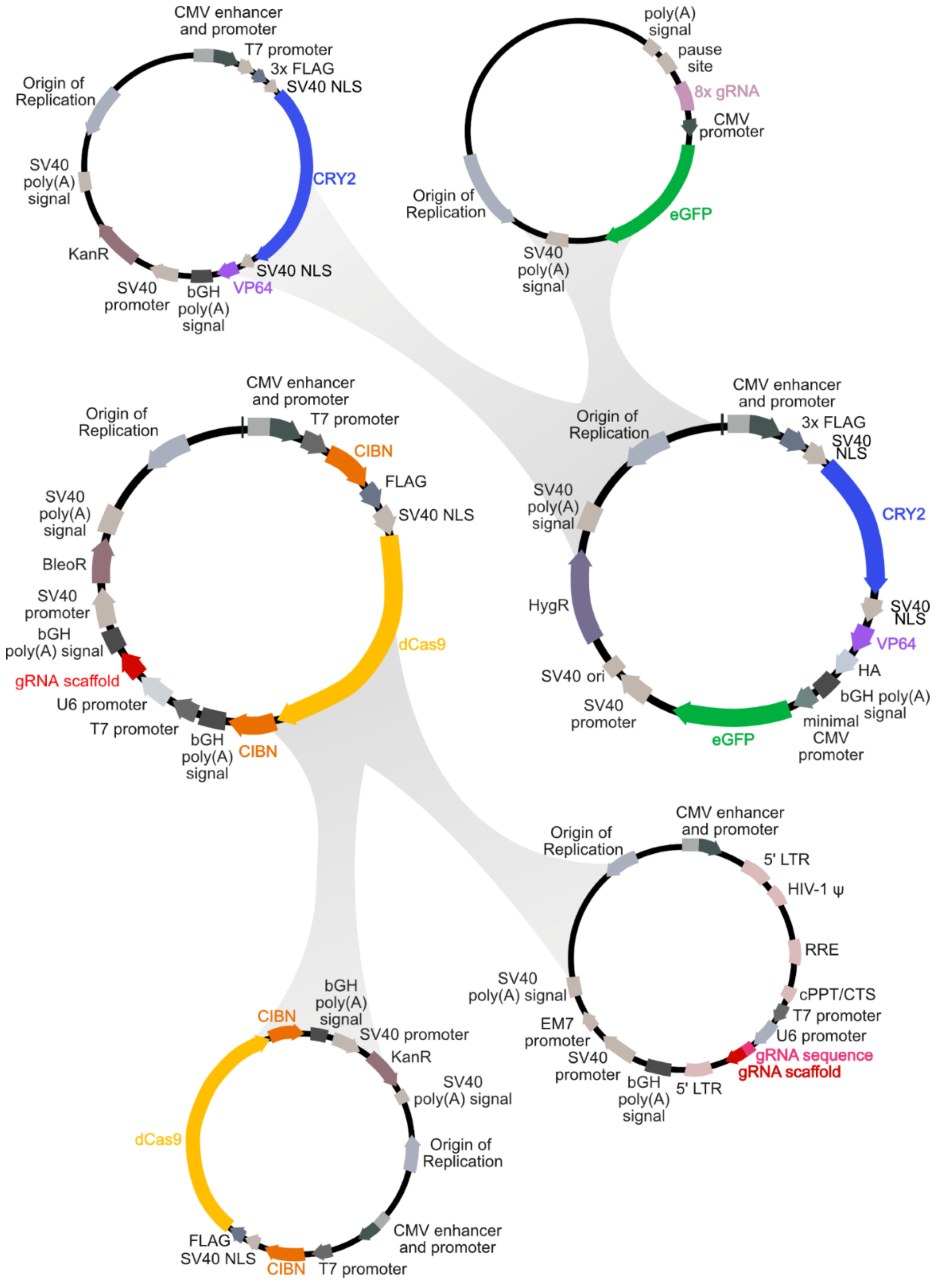

CMV enhancer and promoter
T7 promoter
3x FLAG
SV40 NLS
Origin of Replication
SV40 poly(A) signal
CRY2
KanR
SV40 promoter
bGH poly(A) signal
VP64
SV40 NLS
poly(A) signal
pause site
8x gRNA
CMV promoter
Origin of Replication
eGFP
SV40 poly(A) signal
CMV enhancer and promoter
Origin of Replication
T7 promoter
CIBN
FLAG
SV40 NLS
SV40 poly(A) signal
BleoR
SV40 promoter
bGH poly(A) signal
gRNA scaffold
U6 promoter
T7 promoter
dCas9
bGH poly(A) signal
CIBN
CMV enhancer and promoter
Origin of Replication
3x FLAG
SV40 NLS
SV40 poly(A) signal
CRY2
HygR
SV40 NLS
VP64
SV40 ori
HA
bGH poly(A) signal
SV40 promoter
minimal CMV promoter
eGFP
CMV enhancer and promoter
Origin of Replication
5' LTR
HIV-1 ψ
RRE
SV40 poly(A) signal
cPPT/CTS
T7 promoter
EM7 promoter
U6 promoter
gRNA sequence
SV40 promoter
gRNA scaffold
bGH poly(A) signal
5' LTR
bGH poly(A) signal
CIBN
SV40 promoter
KanR
SV40 poly(A) signal
dCas9
Origin of Replication
CMV enhancer and promoter
FLAG
SV40 NLS
CIBN
T7 promoter

**Figure S1.** Schematic of plasmid design from the original four LACE plasmids (4pLACE) to the optimized two plasmids (2pLACE).

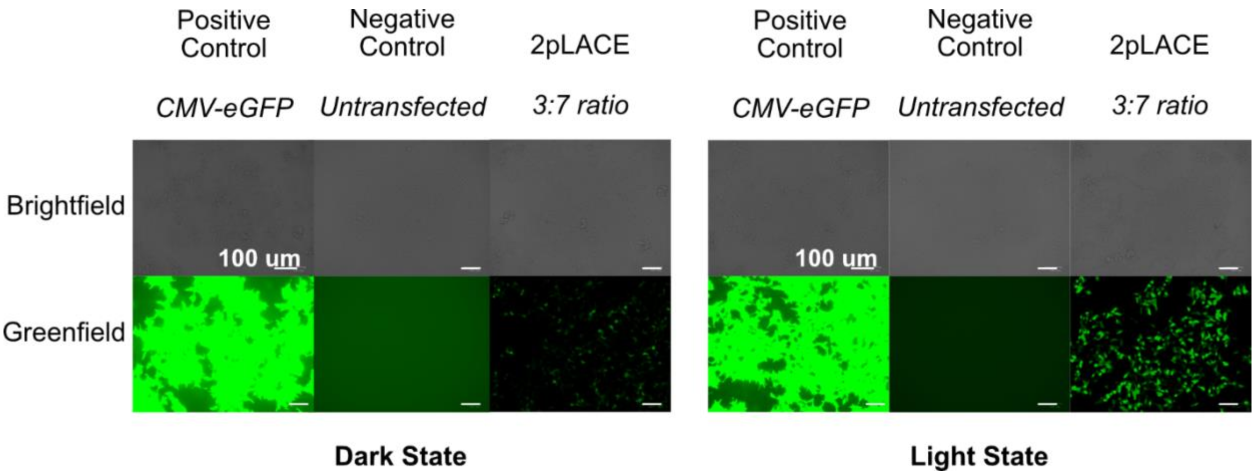


**Figure S2.** eGFP expression of the optimized plasmid ratio for 2pLACE with and without light activation. Images were taken 24 hours post activation.

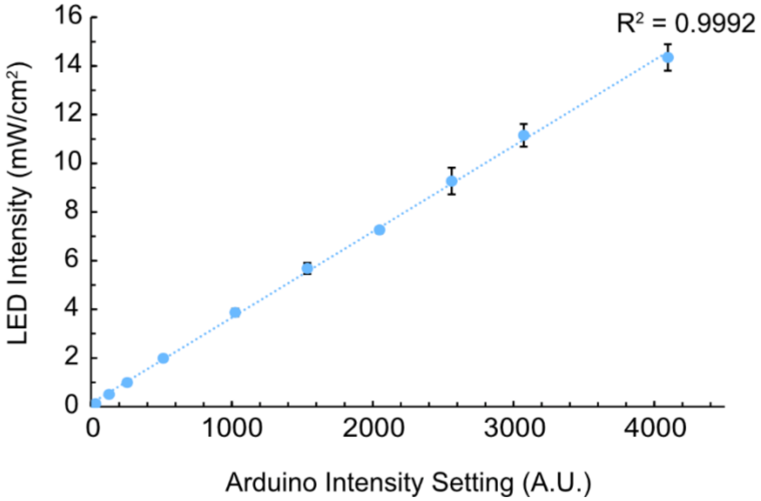


**Figure S3**. Calibration curve of the optoPlate-96. Four LEDs were sampled from the optoPlate-96. Error bars represent standard deviation. Data were fitted with a line and the intercept was fixed to be 0.
